# Frizzled3 inhibits Vangl2-Prickle3 association to establish planar cell polarity in the vertebrate neural plate

**DOI:** 10.1101/2021.05.02.442356

**Authors:** Ilya Chuykin, Keiji Itoh, Kyeongmi Kim, Sergei Y. Sokol

## Abstract

The orientation of epithelial cells in the plane of the tissue, known as planar cell polarity (PCP), is regulated by interactions of asymmetrically localized PCP protein complexes. In the *Xenopus* neural plate, Van Gogh-like2 (Vangl2) and Prickle3 (Pk3) proteins form a complex at the anterior cell boundaries, but how this complex is regulated *in vivo* remains largely unknown. Here we show that Vangl2-Pk3 association is inhibited by Frizzled3 (Fz3), a core PCP protein that is specifically expressed in the neuroectoderm and is essential for the establishment of PCP in this tissue. Proximity biotinylation and crosslinking studies revealed that the Vangl2-Pk3 interaction is suppressed by overexpressed Fz3, but enhanced in Fz3 morphants. In addition, Fz3 induced Vangl2 phosphorylation on T76 and T78, and this phosphorylation was required for Fz3-mediated inhibition of Vangl2-Pk3 complex formation. Consistent with this observation, the complex of Pk3 with nonphosphorylatable Vangl2 was not polarized in the neural plate. These findings provide evidence for *in vivo* regulation of the Vangl2-Pk3 complex formation and localization by a Frizzled receptor.

## INTRODUCTION

Planar cell polarity (PCP) coordinates cell shape, orientation and cell movements that are necessary for normal morphogenesis (Butler & Wallingford, 2017; Devenport, 2014; Goodrich & Strutt, 2011). Initially discovered in *Drosophila*, ‘core PCP’ genes encode the transmembrane proteins Van Gogh (Vang), Frizzled (Fz), Flamingo/Starry Night (Fmi) and the cytosolic components Prickle (Pk) and Dishevelled (Dsh) (Adler, 2012; Peng & Axelrod, 2012). Besides controlling the orientation of epithelial cells, vertebrate PCP gene homologues acquired additional roles in the positioning and functions of subcellular structures, such as basal bodies and cilia, cell and tissue asymmetries, and diverse cell behaviors during development (Davey & Moens, 2017; Gray *et al*, 2011; Sokol, 2015).

A hallmark of PCP in *Drosophila* epithelia is the formation of the core PCP complexes Fmi-Fz-Dsh and Fmi-Vang-Pk that segregate to different cell sides (Peng & Axelrod, 2012). The partitioning of PCP complexes to opposite cell sides is based on their stabilization by intercellular and intracellular feedback interactions (Chen *et al*, 2008; Strutt *et al*, 2011; Tree *et al*, 2002; Wu & Mlodzik, 2008). Vang and Van Gogh-like 2 (Vangl2) can directly associate with Pk *in vitro* (Bastock *et al*, 2003; Chu *et al*, 2016; Jenny *et al*, 2003), but how this complex is regulated in developing vertebrate embryos remains unknown. Wnt5a- and Ror2-dependent phosphorylation of Vangl2 has been proposed to regulate PCP in the developing mouse limb (Gao *et al*, 2011). In *Drosophila*, Vang is also phosphorylated at the conserved sites in response to Fz, but the mechanistic role of this phosphorylation in PCP signaling is not fully understood (Kelly *et al*, 2016).

In the *Xenopus* neural plate, Vangl2 and Pk3 form protein complexes that are enriched at the anterior sides of neuroepithelial cells (Ossipova *et al*, 2015). Importantly, loss-of-function experiments have shown essential roles of Vangl2 in neural tube closure (Darken *et al*, 2002; Goto & Keller, 2002; Greene *et al*, 1998; Jessen *et al*, 2002; Kibar *et al*, 2001; Murdoch *et al*, 2001; Park & Moon, 2002). Fz3 is another core PCP protein that is necessary for neural tube closure in mice (Wang *et al*, 2006). In this study, we use proximity biotinylation and crosslinking approaches (Choi-Rhee *et al*, 2004; Mattson *et al*, 1993; Roux *et al*, 2012) to demonstrate that Fz3 is required for PCP and negatively regulates the association of Vangl2 and Pk3. This function of Fz3 depends on the phosphorylation of the conserved T76 and T78 residues of Vangl2. This phosphorylation is required for the planar polarization of the Vangl2-Pk3 complex in the neural plate. Taken together, our findings reveal the regulation of the Vangl2-Pk3 complex by Fz3-dependent phosphorylation *in vivo*.

## RESULTS

### Fz3 is required for neural plate PCP

To evaluate potential regulation of the Vangl2-Pk3 complex by Frizzled, we studied Fz3 that is expressed in the *Xenopus* neural plate (Shi *et al*, 1998) and has been implicated in mouse neural tube closure (Wang *et al.*, 2006). We first assessed the effect of Fz3 depletion on the localization of Vangl2, which has been shown to accumulate at anterior cell boundaries (Ossipova *et al.*, 2015). Endogenous Vangl2 distribution was analyzed after the immunostaining of neurula stage embryos injected with a previously characterized Fz3 morpholino oligonucleotide (Fz3 MO1)(Deardorff *et al*, 2001) together with GFP as a lineage tracer (Fig. 1A). The anterior enrichment of Vangl2 has been lost in the cells injected with Fz3 MO1 but not in those injected with control MO (Fig. 1B-D). These experiments indicate that Fz3 is essential for the establishment of PCP in the *Xenopus* neural plate.

**Figure 1.**
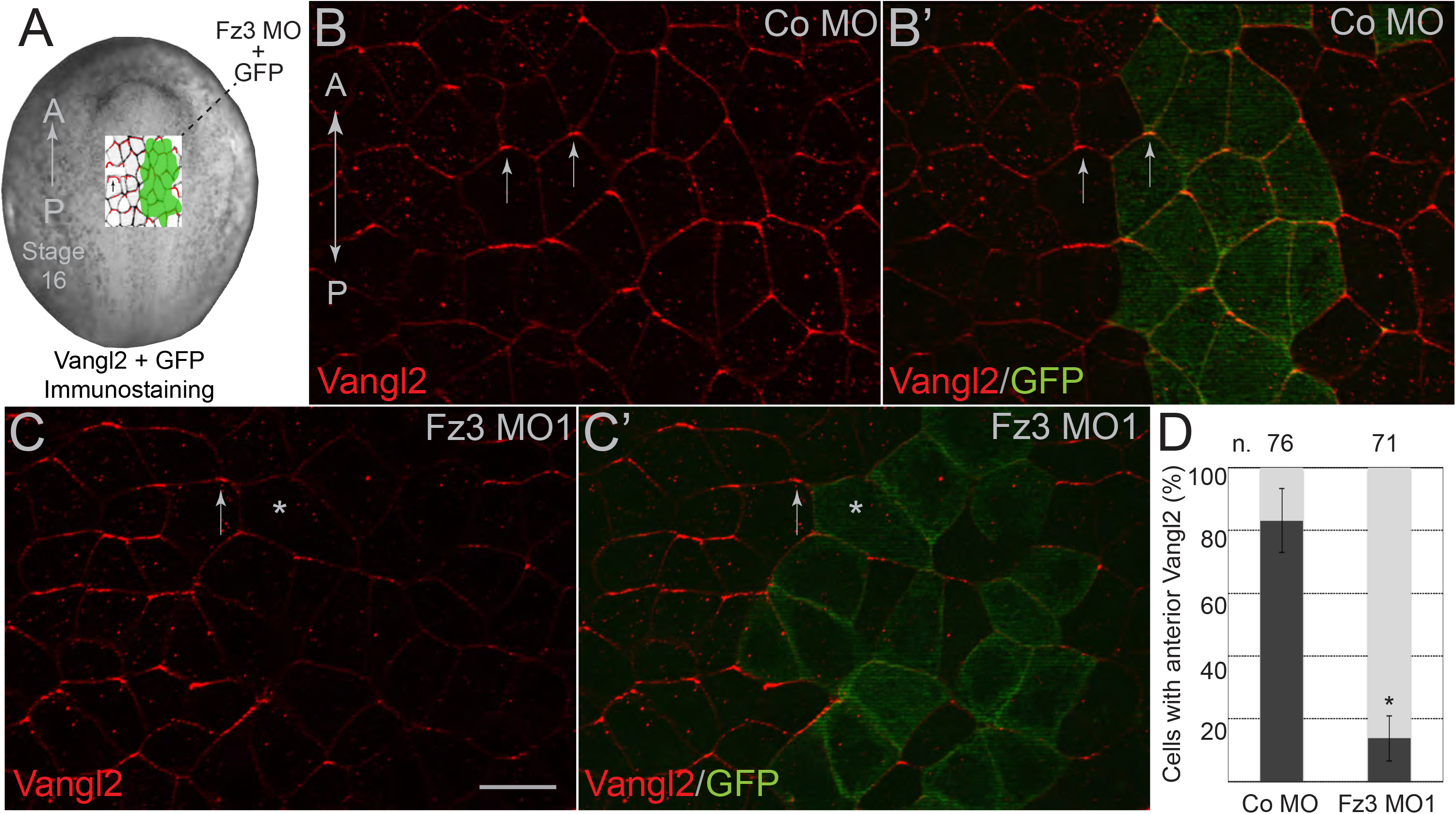
Frizzled3 is required for neural plate planar polarity. (A) Experimental scheme. One dorsal animal blastomere of 16-cell *Xenopus* embryos was co-injected with Fz3 MO1 or control (Co) MO (5 ng each) and GFP RNA (200 pg). Stage 16 embryos were fixed and the neural plates were dissected and stained with anti-Vangl2 and anti-GFP antibodies. (B-C’) Representative *en face* images of neural plates from the injected embryos. Vangl2 (red) is enriched at the anterior cell edges (arrows). GFP is a lineage tracer (green). A-P, anterior-posterior axis. Scale bar, 30 μm. (B, B’) Co MO, (C, C’) Fz3 MO1. Many GFP-positive cells lack Vangl2 polarization (asterisk). (D) Quantification of the mean frequencies ± s. d. for cells with anteriorly enriched Vangl2. Numbers of scored cells (n.) are shown on top of each bar. Three embryos were scored for each group (20-30 cells per embryo). Data represent three different experiments. Statistical significance was determined by the two tailed Student’s *t* test, *, p < 0.05.

### Inhibition of the Vangl2-Pk3 interaction by Fz3

Since the standard immunoprecipitation failed to detect the Vangl2-Pk3 complex in embryos (see below), this protein interaction was monitored by a proximity biotinylation assay. In this approach, we analyzed the biotinylation of exogenous Vangl2 by an N-terminal fragment of the biotin ligase from *Aquifex aeolicus* (Kim *et al*, 2016) that has been fused to Pk3 (BLN-Pk3)(Chuykin *et al*, 2018). Co-expressed Fz3 inhibited Vangl2 biotinylation in a dose-dependent manner, indicating a negative regulation of the Vangl2/Pk3 complex (Fig. 2A). Notably, in the presence of Fz3, Vangl2 protein gel mobility has been reduced, whereas co-expression of Pk3 led to the faster migration of Vangl2 (Fig. 2A, B). These changes in Vangl2 mobility likely indicate phosphorylation, as previously demonstrated for *Drosophila* Vang (Kelly *et al.*, 2016; Strutt *et al*, 2019).

**Figure 2.**
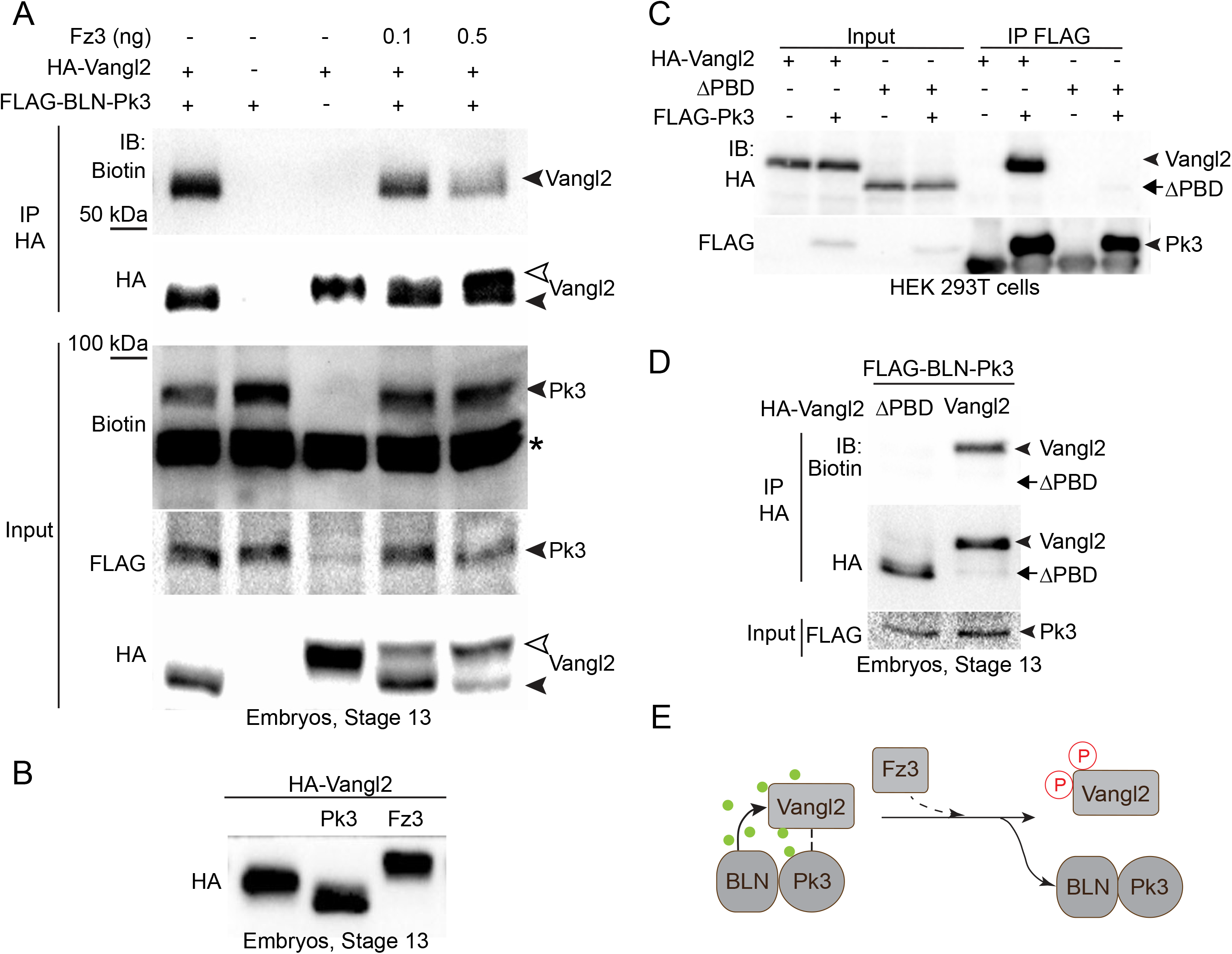
Fz3 reduces the interaction between Vangl2 and Pk3 proteins in *Xenopus* embryos. (A) Vangl2 biotinylation by BLN-Pk3 is decreased in the presence of Fz3. Biotin and mRNAs encoding FLAG-BLN-Pk3 (400 pg), HA-Vangl2 or ΔPBD RNAs (100 pg each) and Fz3-FLAG (100 or 500 pg) as indicated, were injected into the animal region of four- to eight-cell embryos. Protein lysates were immunoprecipitated (IP) with anti-HA antibody from stage 13 *Xenopus* embryos. Proteins were detected by immunoblotting (IB) with anti-biotin, anti-HA and anti-FLAG antibodies as indicated. Black arrowheads point to Vangl2 and Pk3, white arrowhead points to the slower migrating Vangl2 in pulldowns and lysates. Asterisks indicate endogenous proteins labelled by the anti-biotin antibody. (B) Lysates of embryos overexpressing Pk3 and Fz3 show that Pk3 increases Vangl2 mobility, whereas Fz3 reduces it. (C) Physical interaction of HA-Vangl2 and FLAG-Pk3 in transfected 293T cells. HA-Vangl2, but not the construct lacking the presumed Pk3 binding domain (ΔPBD, 298-382 residues), is detected in pulldowns of FLAG-Pk3 from lysates of the transfected HEK293T cells. (D) HA-tagged Vangl2 but not ΔPBD is biotinylated by FLAG-BLN-Pk3. Embryo microinjection details and abbreviations are as in (A). (E) Model. The formation of a complex between Vangl2 and Pk3 results in the biotinylation of Vangl2 by BLN-Pk3, green circles depict biotin. Fz3 causes an upshift of Vangl2 due to phosphorylation (circled red P) and inhibits the Vangl2-Pk3 interaction.

We wanted to confirm that Vangl2 biotinylation truly reflects the physical proximity of the two proteins and generated a Vangl2 deletion construct lacking the presumed Pk3-binding domain (PBD) of Vangl2 (Jenny *et al.*, 2003). As we expected, the Vangl2ΔPBD construct failed to bind Pk3 in transfected HEK293T cells (Fig. 2C) and has not been biotinylated by BLN-Pk3 (Fig. 2D), thereby verifying our approach. These observations indicate that Fz3 inhibits the association of Vangl2 and Pk3 in *Xenopus* embryos (Fig. 2E).

To further verify our conclusion regarding the negative role of Fz3 in the formation of the Vangl2/Pk3 complex, we used immunoprecipitation combined with crosslinking. Whereas the Vangl2-Pk3 complex is readily detectable by immunoprecipitation in cultured cells (Fig. 2C)(Chu *et al.*, 2016), our initial attempts to visualize it in embryos were unsuccessful. We suspected that the Vangl2-Pk3 complex is unstable in a detergent-containing buffer and stabilized it by crosslinking with dithiobis-succinimidyl-propionate (DSP, see Methods) (Fig. 3A). Indeed, Vangl2 was present in Pk3 pulldowns prepared from the crosslinked ectodermal explants, but not in the samples without crosslinking (Fig. 3B). Importantly, significantly lower amounts of Vangl2 coprecipitated with Pk3 from Fz3-expressing explants. Notably, Pk3 preferentially associated with the non-phosphorylated (fast migrating) form of Vangl2. By contrast, both phosphorylated and non-phosphorylated forms of Vangl2 were visible in the cell lysates expressing Fz3 (Fig. 3B). These observations suggest that the PCP complex is modulated by Vangl2 phosphorylation. Taken together, our results validate proximity biotinylation for the analysis of Vangl2-Pk3 association *in vivo* and reveal the negative regulation of this complex by Fz3.

**Figure 3.**
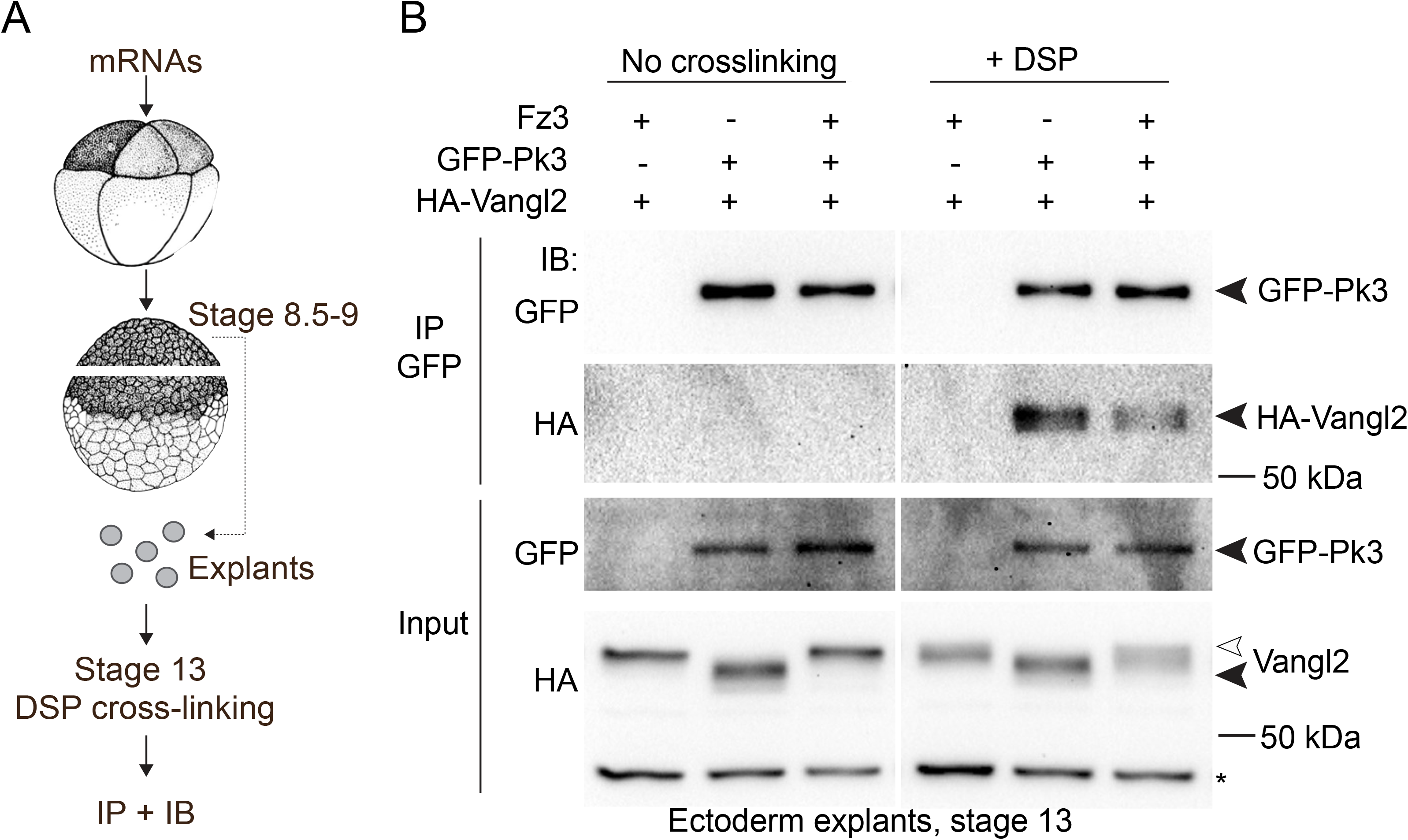
Fz3 decreases the amount of the Vangl2-Pk3 complex *in vivo*. (A) Experimental scheme; eight-cell embryos were injected into each animal blastomere with GFP-Pk3 (200 pg), HA-Vangl2 (50 pg), or Fz3-FLAG (400 pg) mRNA as indicated. Ectoderm explants were dissected at stage 8.5-9, cultured until stage 13, and cross-linked with 2 mM DSP for 30 min. After immunoprecipitation (IP) with GFP-trap beads, protein levels were assessed by immunoblotting (IB) with anti-HA and anti-GFP antibodies. (B) Vangl2 and GFP-Pk3 coprecipitation in control and Fz3-expressing ectoderm explants (right). No binding is detected without cross-linking (left). Black arrowheads point to Vangl2 and Pk3, white arrowhead points to the slower migrating Vangl2 band in the lysates. A nonspecific band reflects protein loading (asterisk).

We next assessed the effect of Fz3 depletion on Vangl2-Pk3 complex formation. Given the dramatic increase in Fz3 transcription during neurulation (Deardorff *et al.*, 2001; Shi *et al.*, 1998), we measured Vangl2 biotinylation at stage 18 to allow sufficient time for protein depletion (Fig. 4A). Fz3 depletion with two different MOs significantly increased Vangl2 biotinylation by BLN-Pk3 (Fig. 4B, C). We also found that levels of endogenous Fz3 were reduced in stage 16-17 Fz3 morphant embryos confirming the efficiency of Fz3 MOs (Fig. 4D, E). These results further support our conclusion that Fz3 inhibits the interaction between Vangl2 and Pk3 *in vivo*.

**Figure 4.**
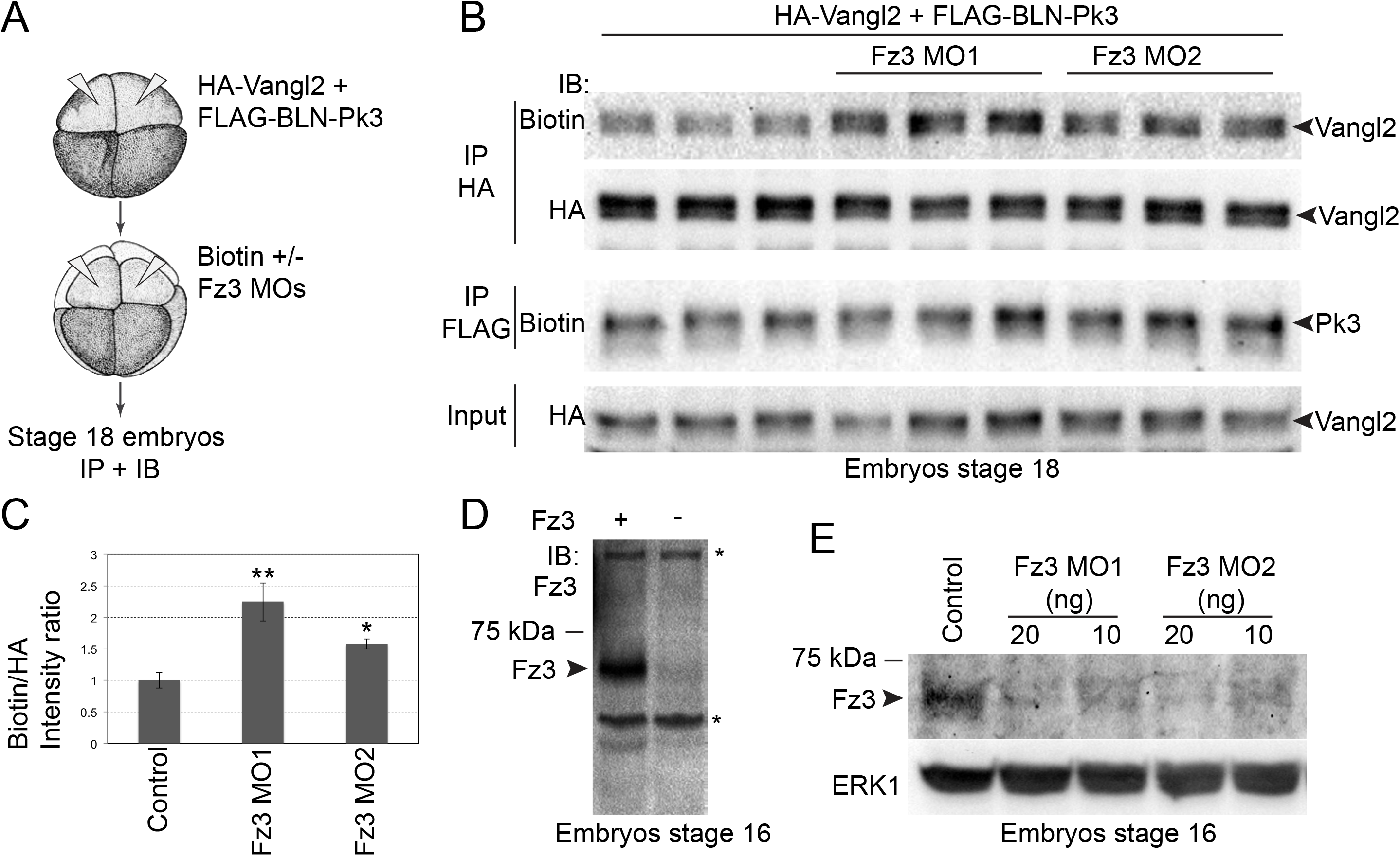
The interaction between Vangl2 and Prickle3 is enhanced in Fz3-depleted embryos. (A) Experimental scheme. Four-to-eight-cell embryos were sequentially injected into dorsal animal blastomeres with BLN-Pk3 and HA-Vangl2 DNAs, 50 pg each, and Fz3 MO1 or Fz3 MO2 with biotin. Embryo lysates were collected at stage 18 for immunoprecipitation (IP) and immunoblotting (IB). (B) Vangl2 and Pk3 biotinylation in control and Fz3 morphant embryos in pull-downs were assessed with anti-biotin antibodies. Total exogenous Vangl2 levels were analyzed by anti-HA antibodies. Due to low abundance and insufficient sensitivity of the detection, the levels and the activity of FLAG-BLN-Pk3 were assessed in the second sequential FLAG pulldown. Biological triplicates were done for each experimental condition with 20 embryos per sample. (C) Band intensity ratios of biotinylated to total Vangl2 levels in pulldowns, evaluated with anti-biotin and anti-HA antibodies, respectively. Statistical significance was determined by the two-tailed Student’s *t*-test, **, p<0.001, *, p<0.05. (D) Anti-Fz3 antibody detects Fz3 (arrowhead) in lysates of stage 16 control embryos or embryos injected with Fz3-FLAG mRNA (25 pg); asterisks indicate non-specific bands. (E) Levels of the endogenous Fz3 protein (arrowhead) in control, Fz3 MO1- and Fz3 MO2-injected stage 16 embryos; ERK1 is a loading control.

### Fz3 induces threonine phosphorylation of Vangl2

Mouse Vangl2 can be phosphorylated at multiple conserved serine/threonine (S/T) residues that belong to two N-terminal clusters (Gao *et al.*, 2011). To identify specific residues that are phosphorylated in response to Fz3, we generated several mutated Xenopus Vangl2 constructs (Fig. 5A), expressed them in the embryo and probed lysates with S82S84-phosphopeptide-specific antibodies (Gao *et al.*, 2011) and anti-phospho-threonine antibodies. Phosphopeptide antibody specificity has been confirmed by lack of staining of the sample containing S82S84>AA (Fig. 5B). The phosphorylation of the S82 and S84 sites did not change in Fz3-expressing ectoderm. By contrast, threonine phosphorylation was induced in several Vangl2 constructs after Fz3 overexpression (Fig. 5B). No change in threonine phosphorylation was observed in the S82S84>AA construct, consistent with S82 and S84 being the founder sites that are necessary for subsequent phosphorylation of other cluster I residues (Gao *et al.*, 2011). Importantly, no increase in phospho-threonine levels was observed in the T76T78>AA construct (Fig. 5C). Based on this evidence, we identify T76 and T78 rather than S82 and S84, as the Vangl2 sites phosphorylated in response to Fz3 signaling.

**Figure 5.**
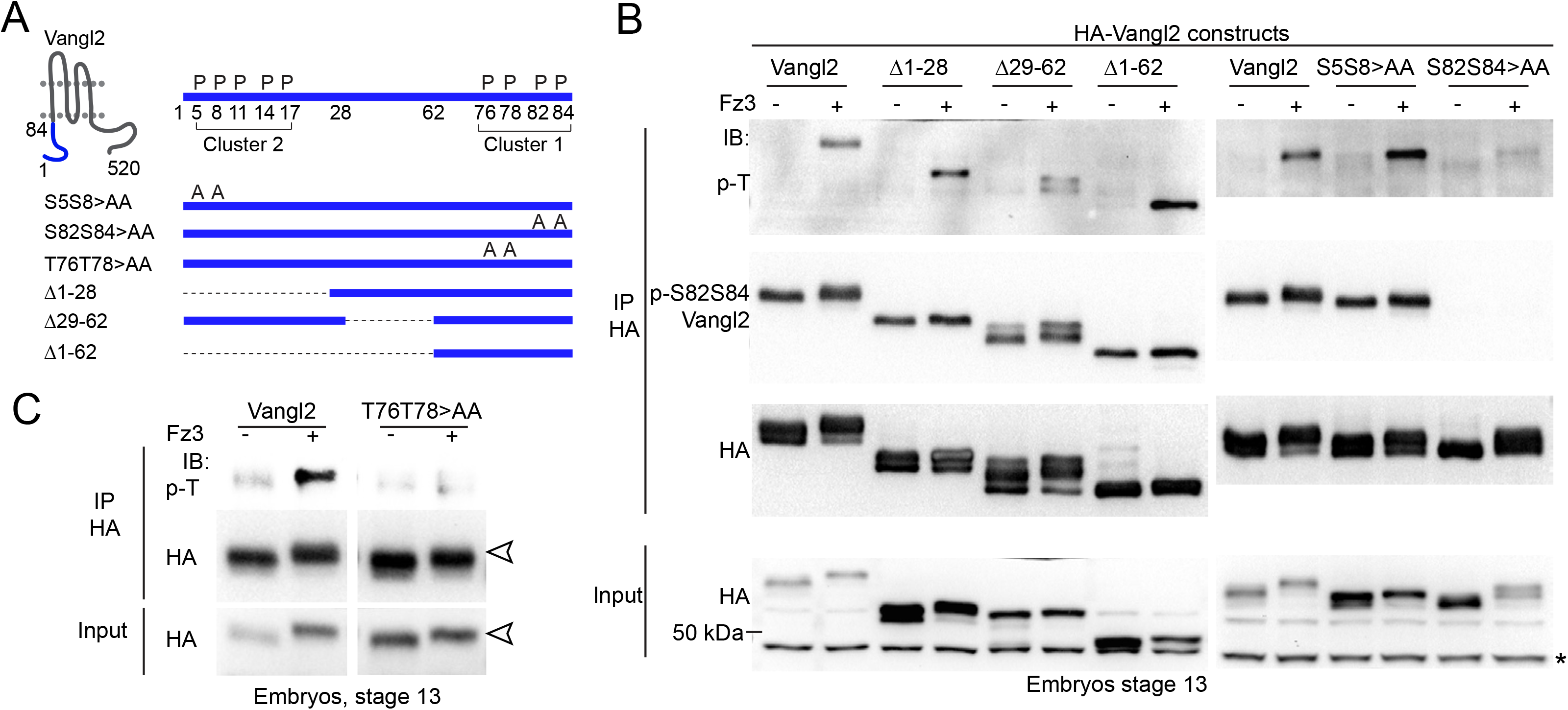
Fz3 induces the phosphorylation of T76 and T78 in Vangl2. (A) Scheme of Vangl2 constructs with the modified cytoplasmic N-terminal region (amino acids 1-84, blue) used to analyze Vangl2 phosphorylation. Phospho-S/T residues in clusters I and II are indicated. (B, C) Immunoprecipitations (IP) with anti-HA antibodies from lysates of embryos expressing Vangl2 constructs with or without Fz3 were immunoblotted (IB) with phospho-specific antibodies as indicated. Anti-HA antibodies show total Vangl2 levels. Asterisk in B marks non-specific band recognized by anti-HA antibody in lysates. (C) Fz3 induces phosphothreonine phosphorylation of the T76T78 sites in Vangl2. Vangl2 upshift is visible in the T76T78>AA mutant (white arrowheads).

### Vangl2 phosphorylation at T76/T78 contributes to PCP by mediating the inhibitory activity of Fz3 on the Vangl2-Pk3 complex

The observed correlation between the reduced Vangl2-Pk3 interaction and the increase in the slow migrating Vangl2 band in Fz3-expressing embryos (Fig. 2B) suggests that Fz3 inhibits Vangl2-Pk3 complex formation by triggering Vangl2 phosphorylation. To address this hypothesis, we tested several Vangl2 constructs with alanine substitutions of the conserved S/T residues (Fig. 6A). We found that the interaction of Pk3 with the T76AT78>AA construct was barely inhibited by Fz3, uncovering roles of T76 and T78 phosphorylation in this process (Fig. 6B). Because the effect of T76T78>AA on Fz3 activity was incomplete, we thought that the S/T residues of cluster II may contribute to the modulation of the PCP complex. This possibility was confirmed after testing additional Vangl2 alanine substitutions. The association of Pk3 with Vangl2-7A was completely resistant to Fz3, whereas the complex containing Vangl2-5A was partially sensitive (Fig. 6A, C). These experiments indicate that the T76T78 phosphorylation is necessary for the inhibitory effect of Fz3 on PCP complex formation.

**Figure 6.**
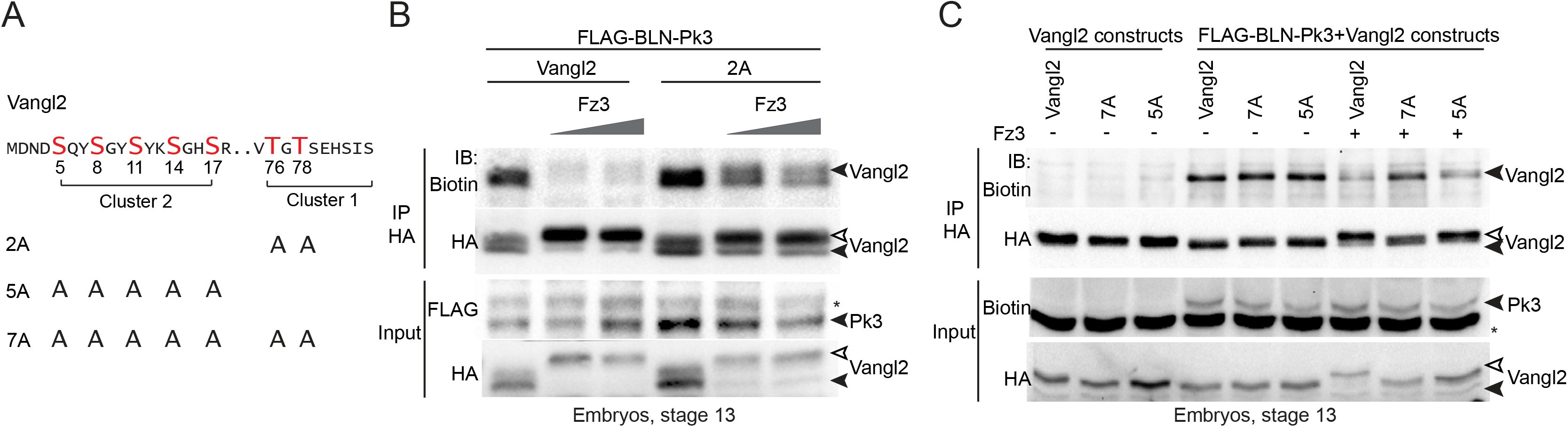
Fz3-induced Vangl2 phosphorylation is necessary for the inhibition of the Vangl2-Pk3 interaction. (A) Scheme of Vangl2 constructs with alanine substitutions. Conserved S/T clusters I and II are indicated. (B, C) Effect of Fz3 on the biotinylation of various Vangl2 constructs by BLN-Pk3. Embryos were injected dorsoanimally with mRNAs encoding Vangl2 constructs (50 pg each), BLN-Pk3, 200 pg, or Fz3-FLAG mRNA, 200 pg or 400 pg (B) and 400 pg (C); and collected at stage 13. (B) Vangl2-2A mutant is partly resistant to the Fz3 inhibitory effect. (C) Vangl2-7A is completely resistant to Fz3 and is not upshifted (black and white arrowheads). Immunoblotting (IB) with anti-biotin, anti-HA and anti-FLAG antibodies was done as indicated.

We next assessed the contribution of Vangl2 phosphorylation to PCP. Vangl2 was compared with Vangl2-7A, which does not dissociate from the Pk3 protein in response to Fz3. Polarized complexes of Pk3 and Vangl2 were readily detectable at anterior cell faces in the neural plate at stage 14, whereas the complexes of Pk3 and Vangl2-7A were more randomly distributed along the cell cortex (Fig. 7A-C). These data are consistent with the hypothesis that Vangl2 phosphorylation by Fz3 is necessary for the anterior enrichment of the Vangl2-Pk3 complex (Fig. 7D).

**Figure 7.**
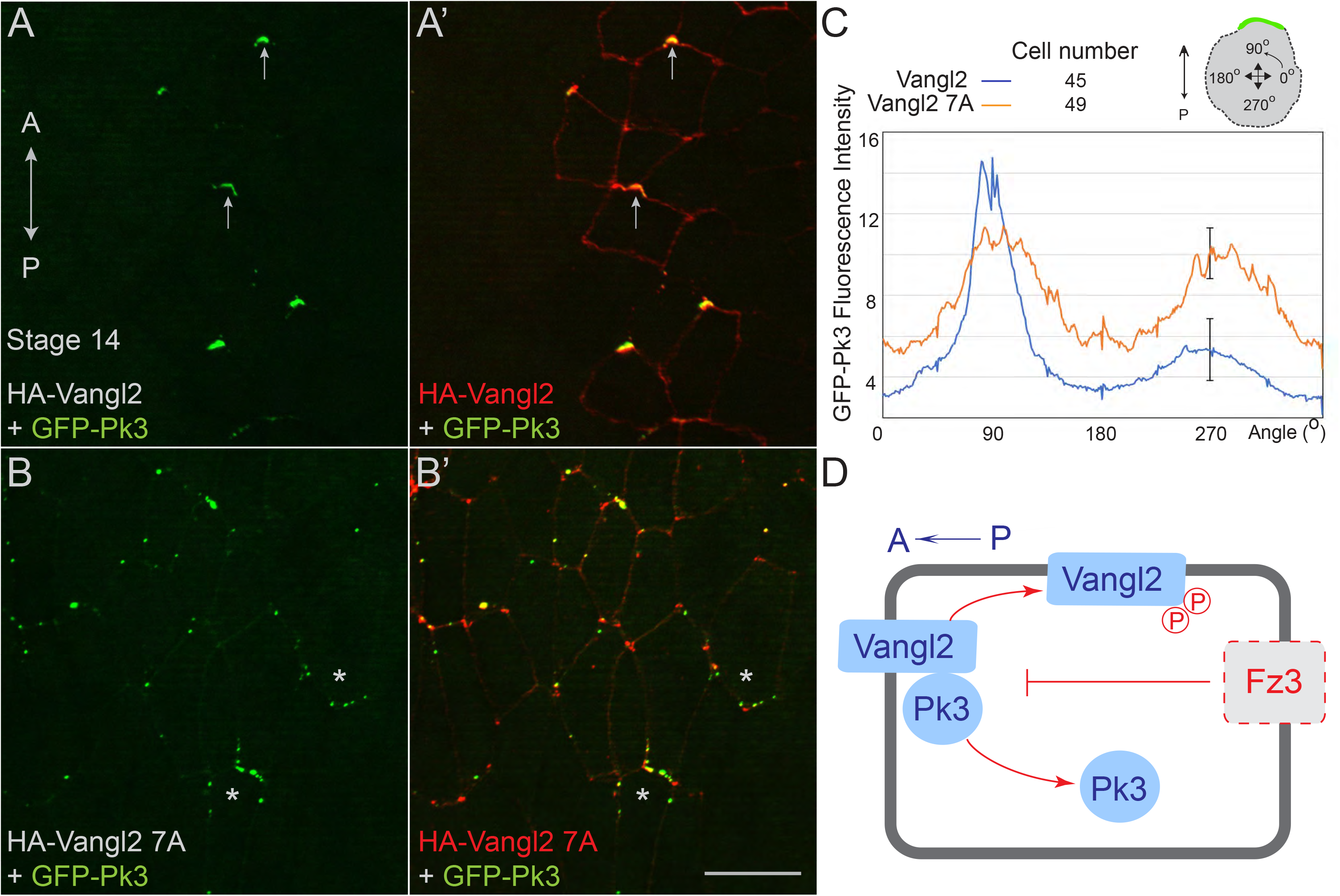
Fz3-induced Vangl2 phosphorylation is required for PCP. (A-C) Two dorsal blastomeres of 16-cell embryos were co-injected with HA-Vangl2 or HA-Vangl2-7A RNA (50 pg) and GFP-Pk3 RNA (200 pg). (A-B’) Representative fixed stage 14 neural plates have been imaged for GFP-Pk3 (direct GFP fluorescence) and immunostained for HA-Vangl2 (A, A’) or HA-Vangl2-7A (B, B’). (A, A’) Polarized anterior PCP complexes are indicated by arrows. (B, B’) Cells lacking anterior GFP-Pk3 are marked by asterisks. Scale bar, 30 μm. Anteroposterior (AP) axis is indicated. (C) Quantification of GFP-Pk3 fluorescence for PCP complexes containing wild-type Vangl2 (blue) and those containing Vangl2-7A (orange) in mosaically expressing cells. Average fluorescence +/- s. d. is shown along the cell circumference for multiple cells as a function of the circular angle from 0 to 360 degrees relative to the AP axis (as shown on the scheme). Numbers of scored cells per each group are indicated. Data are representative of 3 independent experiments. (D) Model. Fz3 contributes to PCP by inhibiting the Vangl2-Pk3 interaction via T76T78 phosphorylation of Vangl2. Presumptive Fz3 localization is shown by dashed line.

## DISCUSSION

Our experiments reveal a role of Fz3 in the inhibition of Vangl2-Pk3 association in the *Xenopus* neural plate. This result supports the hypothesis that negative regulatory feedback between core PCP complexes leads to their segregation to opposite cell sides. In *Drosophila*, Pk has been shown to negatively impact Fz activity (Tree *et al.*, 2002). In vertebrates, Vangl2 can affect the phosphorylation and trafficking of Fz3, although the relevance of this observation to PCP remains to be confirmed (Shafer *et al*, 2011). By contrast, we observe the negative effect of Fz on the Vangl2-Pk complex. While the Vangl2-Pk complex is commonly assumed to be constitutive, we show for the first time its modulation by Fz3. This activity is likely to be allosterically mediated by the phosphorylation of the threonine sites T76 and T78 in Vangl2, at a location that is distinct from the Prickle-binding site of Vangl2. Based on our analysis of nonphosphorylatable Vangl2 constructs, we propose that this activity is required for the establishment of PCP in the neural plate.

If Fz3 antagonizes the Vangl2-Pk3 in a cell-autonomous manner, it should be distributed in a pattern that is complementary to the distribution of the Vangl2-Pk3 aggregates. So far, the presence of the Fz3-containing complex at the posterior sides of neuroepithelial cells has not been documented. Nevertheless, asymmetric localization of Frizzled proteins was reported in other vertebrate tissues, including tracheal and kidney epithelia, inner ear and brain ventricles (Davey *et al*, 2016; Kunimoto *et al*, 2017; Montcouquiol *et al*, 2006; Vladar *et al*, 2012), consistent with a conserved signaling mechanism.

Our experiments identify T76 and T78 as specific amino acid residues that are phosphorylated in response to Fz3 and required for the inhibition of Vangl2-Pk3 association. The substitution of threonines at these sites with alanines together with the additional substitutions of previously identified serines (Gao et al., 2011) disrupts the anterior enrichment of Vangl2-Pk3 complex, supporting a role of Vangl2 phosphorylation in PCP. Although CK1δ and CK1ε kinases have been implicated in the phosphorylation of the conserved S82S84 sites in Vangl2 in the mouse (Gao *et al.*, 2011; Yang *et al*, 2017) and Vang in *Drosophila* (Kelly *et al.*, 2016; Strutt *et al.*, 2019), there may be additional enzymes involved, because the phosphorylation of T76T78 is inducible whereas the phosphorylation of S82S84 does not change in response to Fz3. Further studies are needed to identify a protein kinase that is activated by Fz3 and phosphorylates the T76T78 sites.

How does Fz3 affect the interaction of Vangl2 and Pk3? One possibility is that the phosphorylation prevents Vangl2 oligomerization and clustering that could be affecting the interaction with Prickle (Belotti *et al*, 2012; Yin *et al*, 2012). Alternatively, Fz3 activity may indirectly modulate Vangl2 by affecting another PCP component, such as Ror2 (Gao *et al.*, 2011), Celsr/Fmi (Chen *et al.*, 2008), Dvl (Bastock *et al.*, 2003; Yang *et al.*, 2017) or Par3 (Chuykin *et al.*, 2018).

Although the requirement of PCP molecules in neural tube closure has been established in genetic studies (Nikolopoulou *et al*, 2017), how they operate during neurulation is still unclear. Notably, we observed that nonphosphorylatable Vangl2 mutants interfere with neural tube closure. PCP complexes have been hypothesized to trigger actomyosin contractility and cell junction shrinking (McGreevy *et al*, 2015; Nishimura *et al*, 2012), but the direct data are still missing. One possibility is that core PCP proteins interact with and modulate actin-remodeling proteins, as has been proposed for the actin-binding protein Shroom3 (Hildebrand & Soriano, 1999; McGreevy *et al.*, 2015; Ossipova *et al*, 2014). Conversely, PCP proteins may themselves be regulated by actomyosin-dependent forces (Chien *et al*, 2015; Mahaffey *et al*, 2013; Newman-Smith *et al*, 2015; Ossipova *et al.*, 2015). These mechanisms are consistent with the genetic evidence that Rock/Myosin II signaling is required for fly PCP (Strutt *et al*, 1997; Winter *et al*, 2001). Future studies should address the possible connection between the Fz3-dependent Vangl2 phosphorylation that we described and cell movements during neurulation.

## MATERIALS AND METHODS

### Plasmids, mRNA synthesis and morpholino oligonucleotides

Plasmids encoding *Xenopus* GFP-, FLAG- and FLAG-BLN-tagged Pk3, HA-Vangl2 and Fz3-FLAG were described previously (Chu *et al.*, 2016; Chuykin *et al.*, 2018; Shi *et al.*, 1998). HA-tagged deletion mutants of Vangl2 Δ1-28, Δ29-62, Δ1-62, Δ298-382 and Vangl2 S/T>A substitutions were generated by site-directed mutagenesis or sub-cloning of gBlock fragments (Integrated DNA Technologies) in pCS2. Capped mRNAs were synthesized using mMessage mMachine kit (Ambion, Austin, TX). For depletion studies, the following MOs were purchased from Gene Tools (Philomath, OR): Fz3 MO1: 5’-CGCAAAGCCACATGCACCTCTTGAA-3’ (Deardorff *et al.*, 2001), Fz3 MO2 5’-CTTGCGTCCAAAGCACCAATTGCTC-3’ and control MO (Co MO) 5’-AGCGTTTCAGGCCGATCTCTCAGTC-3’.

### *Xenopus* embryo culture and microinjections

*In vitro* fertilized eggs were obtained from *Xenopus laevis* and cultured in 0.1x Marc’s Modified Ringer’s solution (MMR) as described previously (Newport & Kirschner, 1982). Staging was according to (Nieuwkoop & Faber, 1967). For microinjections, 4-16-cell embryos were transferred to 3 % Ficoll (GE Healthcare) in 0.6x MMR and injected with 5-10 nl of a solution containing mRNAs and/or MOs. For mosaic expression of PCP complexes, embryos were injected into two dorsal blastomeres of sixteen cell embryos. Amounts of injected mRNAs have been optimized in the preliminary dose-response experiments and are indicated in figure legends.

### Cell culture, transfection and immunoprecipitation

HEK293T cells (ATCC) were maintained in Dulbecco’s modified Eagles medium (Corning) with 10 % fetal bovine serum (Sigma) and penicillin/streptomycin (Sigma). Cells growing at 50-70 % confluence were transiently transfected using linear polyethylenimine (M.W. 25000, Polysciences) as described (Chuykin *et al.*, 2018). For transfections, each 60-mm well received 1 μg of plasmids encoding FLAG-Pk3, HA-Vangl2 WT or HA-Vangl2Δ298-382 constructs in pCS2. Vector DNA (pCS2) was added to plasmid DNA mixture to reach the total amount of 3 μg. For immunoprecipitation, cells were harvested 2 days after transfections in the buffer containing 50 mM Tris-HCl pH 7.5, 150 mM NaCl, 1 mM EDTA and 1 % Triton-X100. Lysates were incubated for several hours or overnight with 5 μl of M2 agarose beads (Sigma) at 4°C on a rotating shaker. The beads were washed twice with the lysis buffer and the samples were heated at 95°C in the sample buffer for separation in SDS-PAGE.

### Immunostaining, fluorescent protein detection, imaging and quantification

To detect PCP complexes in the neural plate, embryos were collected at stage 14 and the vitelline membrane was removed manually. GFP-Pk3 mRNA was co-expressed with Vangl2 mRNA at the previously established doses (as indicated in the figure legends) that have no effect on normal development. Embryos were fixed in MEMFA solution for 40 min, washed in 0.3 % Triton-X-100 in PBS, and neural plate explants were dissected, immunostained with anti-HA antibody (1:100, 12CA5 hybridoma supernatant) to detect HA-Vangl2, imaged and scored. For detection of endogenous Vangl2, embryos were collected at stage 16, fixed in 2 % TCA for 30 min, and the immunostaining was done with the rabbit polyclonal anti-Vangl2 antibody (1:100) (Ossipova *et al.*, 2015). For the tracing of Fz3-depleted cells, embryos were co-injected with MOs and GFP mRNA and co-immunostained with anti-Vangl2 and with mouse monoclonal anti-GFP antibody (GFP-B2, Santa Cruz). Secondary antibodies were against mouse or rabbit IgG conjugated to Cy2 or Cy3 (1:500, Jackson ImmunoResearch). GFP-Pk3 was detected by epifluorescence. Explants were mounted in the Vectashield mounting medium (Vector). Standard specificity controls were performed to confirm the lack of cross-reactivity and no staining without primary antibodies.

Images that are representative of at least 10 different fields were captured using a Zeiss AxioImager microscope with the Apotome attachment. The data shown are from two to five independent experiments with 5-15 embryos per group. To quantify the effect of Fz3 depletion on PCP, cells with anteriorly enriched Vangl2 were scored for three embryos per group, 20-30 cells per embryo, after the embryos have been injected with Fz3 MO or Co MO with GFP as a tracer. Statistical significance was determined by the two tailed Student’s T test (Excel). The quantification of planar polarity of the of GFP-Pk3/HA-Vangl2 complexes (Vangl2 versus Vangl2-7A constructs) was done using the Fiji software with Azimuthal Average plugin https://imagej.nih.gov/ij/plugins/azimuthal-average.html. Mosaic cells were selected using the polygon tool in Fiji and GFP-Pk3 fluorescence intensity was quantified along the radius in cells expressing PCP complexes, the calculated means were plotted as a graph (Microsoft Excel). Data represent 3 independent experiments and were verified blindly.

### Proximity biotinylation in *Xenopus* embryos and immunoblotting

For proximity biotinylation, embryos were injected into the animal pole of 4-8-cell embryos with 10 nl of the solution containing 0.8 mM of biotin and mRNAs encoding FLAG-BLN-Pk3, 200 or 400 pg, HA-Vangl2, 50 pg, and Fz3-FLAG, 200 to 700 pg. Embryos were lysed and the protein biotinylation was assessed in embryo lysates and pulldowns with mouse anti-HA (12A5) and anti-FLAG (M2) antibodies. Stage is indicated in the Figure legends. Proteins were detected by the immunoblotting with goat anti-biotin-HRP antibodies (Cell Signaling) or goat anti-biotin antibody (Pierce), anti-FLAG (M2 Sigma), rabbit anti-HA (Bethyl) and mouse monoclonal anti-*Xenopus* Fz3 antibodies (a gift of Peter Klein)(Deardorff *et al.*, 2001).

### Analysis of the Vangl2-Pk3 binding in crosslinked ectoderm explants

Four- to eight-cell embryos were injected into all four animal blastomeres with mRNAs encoding GFP-Pk3, 200 pg, HA-Vangl2, 50 pg, with or without Fz3-FLAG, 400 pg. Ectoderm explants were isolated at stage 8.5-9 and cultured until stage 13. 30 explants were collected in 50 μl of 0.6× MMR solution and treated with 2 mM dithiobis-succinimidyl-propionate (DSP)(ThermoFisher) (Mattson *et al.*, 1993) for 30 min, lysed and the pull down of GFP-Pk3 was performed with GFP-Trap (Chromotek) followed by immunoblotting.

## Acknowledgement

We thank Peter Klein and De-Li Shi for *Xenopus* Fz3 plasmids and monoclonal antibodies specific for Fz3, Chih-Wen Chu for Fz3 MO2, Yingzi Yang for pS82S84 phosphopeptide-specific antibodies, Shilpa Dilip Kumar for advice on quantification. We also thank Miho Matsuda for comments on the manuscript and members of the Sokol laboratory for discussions. This study has been supported by National Institutes of Health grants GM122492 and NS100759 to SYS.

